# xCell 2.0: Robust Algorithm for cell type Proportion Estimation Predicts Response to Immune Checkpoint Blockade

**DOI:** 10.1101/2024.09.06.611424

**Authors:** Almog Angel, Loai Naom, Shir Nabet-Levy, Dvir Aran

## Abstract

**Background:** Accurate estimation of cell type proportions from bulk gene expression data is essential for understanding the cellular heterogeneity underlying complex tissues and diseases. Here, we introduce xCell 2.0, an advanced version of the xCell algorithm, featuring a training function that permits the utilization of any reference dataset. xCell 2.0 generates cell type gene signatures using an improved methodology, including automated handling of cell type dependencies and more robust signature generation.

**Methods:** We benchmarked xCell 2.0 against ten popular deconvolution methods using nine human and mouse reference sets and 26 validation datasets, encompassing 1,749 samples and 67 cell types. Additionally, we validated xCell 2.0 using the independent Deconvolution DREAM Challenge dataset. As an applicative test case, we curated pan-cancer data of 2,007 patients pre-treated with immune checkpoint blockade (ICB). Features of the tumor microenvironment (TME) were generated using xCell 2.0 and other methods and fed into a LightGBM model using nested cross-validation to obtain robust predictions of ICB response.

**Results:** Benchmarking results showed that xCell 2.0 outperformed all other tested methods across distinct reference datasets, demonstrating superior accuracy and consistency across diverse biological contexts. xCell 2.0 also showed the best performance in minimizing spillover effects between related cell types. In the ICB response prediction task, xCell 2.0-derived TME features significantly improved prediction accuracy compared to models using only cancer type and treatment information, and outperformed other deconvolution methods and established ICB prediction scores.

**Conclusions:** xCell 2.0 is a versatile and robust tool for cell type deconvolution that maintains high performance across various reference types and biological contexts. It is available both via a web application and as a Bioconductor-compatible package, equipped with a large collection of pre-trained cell type signatures for human and mouse research. The improved prediction of ICB responses highlights the potential of xCell 2.0 to advance precision medicine in cancer and other diseases.

## INTRODUCTION

Cellular deconvolution of bulk gene expression data is a powerful tool for uncovering the cellular heterogeneity underlying complex tissues and diseases [1, 2]. While single-cell RNA sequencing (scRNAseq) provides unprecedented resolution of cellular diversity, it is still prohibitively expensive and has limited public data availability [3, 4]. Moreover, scRNA-seq experiments are often performed on freshly isolated cells and therefore not retrospective, making it difficult to use these data for predicting response to treatments [5, 6]. Therefore, there is a continued need for the analysis of bulk gene expression data to gain insights into cellular composition and function [7].

We previously developed xCell, a gene signature-based algorithm that estimates the relative abundance of different cell types in bulk gene expression data [8]. The original version of xCell was pre-trained using reference gene expression datasets and could not be used with custom-made references, limiting its usability for specific tissue types or experimental conditions. For example, one important application of cellular deconvolution is to dissect the complex cellular landscape of the tumor microenvironment (TME) [9, 10]. Since the TME is composed of cell types that are not found in blood, it is important to perform such analyses using TME-dedicated references [11, 12]. While xCell gained popularity, due to its high accuracy and ease of use, it is still important to develop a more flexible version of xCell that can be trained with any given reference dataset.

Here, we present xCell 2.0, an updated version of the xCell algorithm that incorporates a training function and improved gene signature generation process, allowing users to use any given reference dataset. Key changes include automated handling of cell type dependencies and more robust signature generation. Significantly, the versatility of xCell 2.0 enables its application in various scenarios including studies on different diseases, tissues, and even in different organisms such as mice. Moreover, we provide comprehensive pre-trained references within xCell 2.0, covering a diverse range of cell types from various tissues, organs and organisms. We benchmarked xCell 2.0 on 24 human and 2 mouse datasets and compared it to other popular methods, demonstrating that it consistently excels regardless of the training reference being used. Furthermore, we applied xCell 2.0 to a curated dataset of bulk RNA-seq data from 2,007 cancer patients prior to treatment with Immune checkpoint blockade (ICB) from different cancer types. Our results demonstrate that xCell 2.0 can improve the prediction of patient response to ICB therapy, a critical challenge in cancer treatment, highlighting its potential for advancing precision medicine in cancer and other diseases. These findings highlight the potential of xCell 2.0 as a powerful tool for unraveling cellular heterogeneity and guiding personalized therapeutic strategies across a wide range of biomedical applications.

## RESULTS

### Generalized reference-based cell type enrichment

xCell 2.0 introduces a pipeline for generating custom reference objects that can be used for cell type enrichment analysis (Figure 1A). This pipeline, which enhances the applicability of the method to diverse tissue types and experimental conditions, begins by obtaining a reference gene expression dataset of pure cell types, which can be derived from microarray, bulk RNA-seq, or scRNA-Seq data. Next, it generates cell type gene signatures using a similar approach to the original xCell method. Finally, xCell 2.0 generates in-silico simulations to learn the parameters to transform the enrichment scores to linear scores and correct for spillover. These are performed in a similar manner to the original xCell, but using automatic identification of control cell types to avoid manual intervention. The result of this process is a custom reference object that can be used for cell type enrichment analysis of any bulk gene expression datasets.

**Figure 1.**
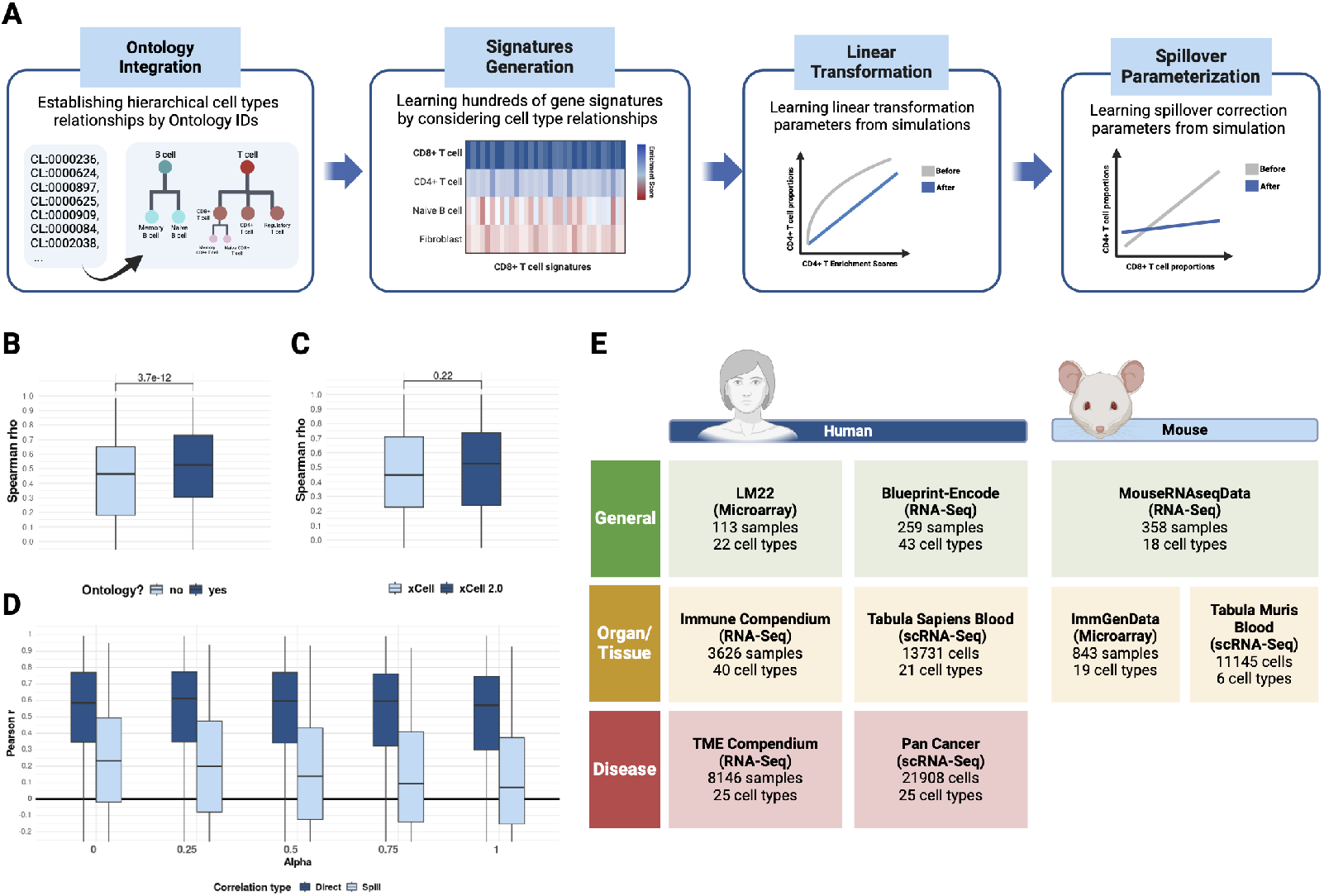
xCell 2.0 pipeline and performance evaluation. **A**. The xCell 2.0 pipeline consists of four main steps: 1) Ontology integration to identify cell type dependencies using standardized Cell Ontology (CL) IDs. 2) Generation of hundreds of gene signatures for each cell type using various thresholds of gene uniqueness. 3) Creation of in-silico simulations to learn parameters for linear transformation of enrichment scores. 4) Spillover parameterization to correct for effects between related cell types. **B**. Accounting for cell type dependencies significantly improves overall signature performance. The graph shows the performance comparison between xCell 2.0 with and without ontology integration. **C**. Evaluation of xCell 2.0’s modified threshold criteria for gene inclusion in signatures. Results show consistent performance with a slight improvement over the original xCell version when tested on the Blueprint-Encode reference. **D**. Effect of spillover correction strength (*α*) on direct and spill correlations. The graph demonstrates that direct correlation remains stable while spill correlation decreases with increasing *α* values. **E**. Overview of pre-trained reference datasets available in xCell 2.0. The figure shows multiple human and mouse reference objects, including bulk RNA-seq, scRNA-seq, and microarray data sources, covering various tissues, immune cell types, and cancer contexts.

We now describe in more detail the four steps, focusing on the changes made in the new version. In the first step, we enhance xCell 2.0 versatility by improving the handling of cell type dependencies caused by lineage relationships. The original xCell required manual identification of these dependencies to ensure that closely related cell types were not directly compared during the signature generation process (e.g., T cells and CD4+ T cells would not be compared to avoid lineage-related biases). However, when dealing with custom references that may contain many cell types, this manual identification process is labor-intensive and requires substantial domain expertise. To overcome this limitation, xCell 2.0 introduces ontological integration. In this step, we automate the identification of lineage relationships among cell types using ontology IDs. xCell 2.0 extracts cell type lineage information directly from the standardized Cell Ontology (CL) [13], enabling the entire pipeline to account for cell type dependencies automatically. Notably, many competing methods do not handle these dependencies at all, requiring users to manually avoid any dependencies in the reference matrix when using deconvolution-based approaches. Using our benchmark validation datasets (described in the next result section), we demonstrate that accounting for cell type dependencies significantly improves overall signature performance (Figure 1B).

Next, xCell 2.0 generates cell type signatures in a similar approach as the original xCell where hundreds of signatures for each cell type are generated using various predefined thresholds. These include thresholds for different percentiles of gene expression, the difference in expression between the cell type of interest and others, the minimum and maximum number of genes per signature, and more. In xCell 2.0 we modified the threshold criteria for determining gene inclusion into signatures. Specifically, the original approach of considering only genes that passed these threshold criteria against the top three other cell types is not feasible when dealing with custom references with an unknown number of cell types. To accommodate the variability in the number of cell types in custom references, we implemented a threshold-based approach of at least 50% of the cell types in the reference. This threshold was chosen as a balance between stringency and flexibility, allowing for robust signature generation across diverse reference datasets. We evaluated this change in the core of the xCell signature generation algorithm using signatures trained on the Blueprint-Encode reference in both xCell and xCell 2.0 on a set of validation datasets. Results show that the overall performance of xCell 2.0 signatures remains consistent with a slight, non-significant, improvement over the first version (Figure 1C).

In the last two steps, we use in-silico simulated cell type mixtures to learn parameters that model the linear relationship between signatures’ enrichment scores and cell type proportions. In xCell 2.0, the control cell type for the mixtures is chosen automatically as the most distinct cell type according to gene expression correlation, from the cell type of interest.

We use the transformed enrichment scores of each cell type in controlled mixtures to measure the spillover effect of those signatures compared to all other cell types. This results in a spillover matrix that reflects the pairwise spillover between all cell types (except cell types with lineage dependencies). Using the spillover correction strength (*α*), one can control for the strength of the spillover effect correction.

This control is crucial as it allows users to balance between correcting for genuine spillover effects and potentially over-correcting, which could introduce new biases. To estimate the effect of *α*, we measured the Pearson correlation between each cell type estimated proportion and the corresponding ground truth proportion (referred to as direct correlation) and again with the ground truth proportion of the most similar cell type in the mixture (referred to as spill correlation) by using different values for *α* (Figure 1D). The results show that the direct correlation remains stable and scarcely affected by the spillover correlation adjustment while the spill correlation decreases as controlling for a higher strength of spillover correlation using *α*.

To facilitate the use of xCell 2.0, we have curated and pre-trained multiple human and mouse reference datasets [14–22], providing extensive ready-to-use objects for a variety of research fields (Figure 1E and Supplementary Table 1). These references encompass a wide range of cell types from various tissues, including comprehensive immune cell compendiums and pan-cancer datasets. We generated reference objects based on both bulk transcriptome data (including microarray and RNA-seq) and single-cell RNA-seq datasets. These pre-trained references cover diverse biological contexts, from specific tissue types to more general applications, ensuring broad applicability across different research areas. All the pre-trained objects are available at https://dviraran.github.io/xCell2refs. We also invite the scientific community to train and share their own custom reference objects.

In summary, xCell 2.0 introduces several key improvements, including automated handling of cell type dependencies, more robust signature generation, and automatically simulating mixtures for parameter learning, all of which contribute to its enhanced performance and applicability across diverse datasets. Furthermore, xCell 2.0 is ready for immediate use with a comprehensive set of pre-trained references covering various human and mouse tissues, immune cell types, and cancer contexts.

### Comparative evaluation of xCell 2.0 performance

We conducted a comprehensive evaluation of the performance of xCell 2.0. Our benchmark strategy utilized nine distinct reference objects (Figure 2A): two human and two mouse general references not associated with specific tissues or diseases, two human and one mouse blood-specific references, and two human TME references. These references were paired with 26 validation datasets (24 human and 2 mouse), each containing bulk gene expression data and corresponding ground truth cell type proportions measured using various validation techniques, primarily cytometry. In total, we analyzed 85 combinations of reference-validation datasets, encompassing 1,749 samples and 67 cell types (Supplementary Table 2).

**Figure 2.**
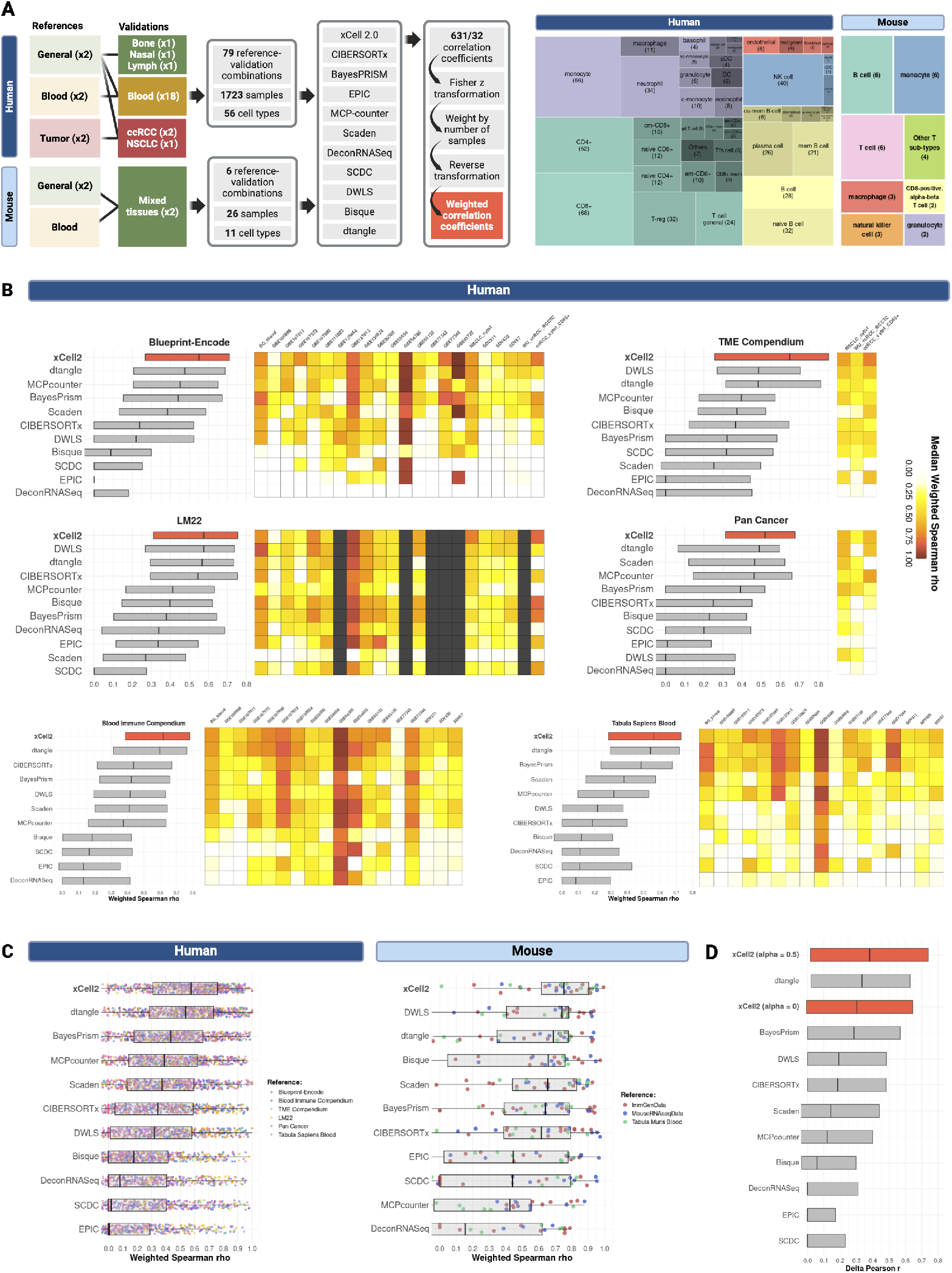
Benchmarking xCell 2.0. **A**. Overview of the benchmarking approach. Nine human and mouse reference datasets were paired with 27 validation datasets spanning various tissue types. The benchmarking process involved 79 reference-validation combinations, encompassing 1,723 samples and 56 cell types for human data (right treemap), and 6 reference-validation combinations with 26 samples and 11 cell types for mouse data (right treemap). Spearman correlation coefficients were weighted to ensure fair comparison across datasets of different sizes. **B**. Performance comparison of xCell 2.0 with ten other deconvolution methods across six human reference datasets. Boxplots show the weighted Spearman coefficients and are ordered by their median. xCell 2.0 outperformed the other methods in all six references used in this study. **C**. Weighted Spearman coefficients of all cell type validations across all references for human (left) and mouse (right) benchmarking sets. Each dot represents a correlation between a cell type estimation by one of the 11 methods and its ground truth with all the different relevant references. **D**. Evaluation of spillover effects. Using a spillover correction strength (*α*) of 0.5, xCell 2.0 demonstrated superior ability in minimizing spillover effects between related cell types compared to other methods.

To assess performance, we calculated both Spearman and Pearson correlation coefficients between the predicted and ground-truth proportions for each method. However, we chose to use Spearman correlation as our primary metric, as it is more relevant for cell type estimation problems. Spearman correlation better captures the ability to distinguish between small differences in cell type proportions, which are often more critical in downstream analyses. Pearson correlation, in contrast, can be dominated by large differences in cell type proportions, potentially obscuring a method’s performance in distinguishing subtle variations. To account for varying sample sizes across validation datasets, we applied a weighting procedure. First, we transformed the correlation coefficients to z-scores using Fisher’s transformation. We then weighted these z-scores by the natural logarithm of the number of samples in each dataset. Finally, we applied the inverse Fisher transformation to convert the weighted z-scores back to correlation coefficients. This approach ensures a fair comparison across datasets of different sizes while maintaining the interpretability of correlation coefficients (Figure 2A). All figures in the main text present Spearman correlations, while Pearson correlations are provided in the supplementary materials for completeness.

We extended our benchmark strategy to evaluate ten widely-used deconvolution methods: BayesPrism [23], Bisque [24], CIBERSORTx [1], DeconRNAseq [25], dtangle [26], DWLS [27], EPIC [28], MCP-counter [29], Scaden [30], and SCDC [31]. The comparative analysis revealed that xCell 2.0 outperformed all other methods in all six human references (Figure 2B and Supplementary Figure 1). Combined for all the references together, the weighted median correlation coefficient for xCell 2.0 was 0.58, compared to 0.54 for dtangle in second place and 0.43 for BayesPrism in third place (Figure 2C). For the mouse datasets, xCell 2.0 maintained its superior performance, ranking first with a weighted median correlation coefficient of 0.75, followed closely by DWLS at 0.74 and dtangle at 0.69 (Figure 2C and Supplementary Figure 2). This consistent top performance across both human and mouse datasets demonstrates xCell 2.0’s versatility and robustness in deconvoluting gene expression data from various species and tissue types.

To further validate the performance of xCell 2.0, we leveraged the Tumor Deconvolution DREAM Challenge, a community-driven benchmarking framework specifically designed to evaluate deconvolution methods [32]. Utilizing this independent and comprehensive framework allowed us to robustly compare xCell 2.0 against a wide range of state-of-the-art methods, thereby strengthening the validity of our findings. This analysis was conducted across both coarse and fine-grained cell type annotations, enabling a detailed assessment of the accuracy and robustness of xCell 2.0 in capturing cell heterogeneity. In the analysis of coarse-annotated cell types, xCell 2.0 demonstrated strong performance, achieving second place out of 23 methods with a Spearman correlation of 0.776. This performance was only surpassed by CIBERSORTx (rho = 0.831) and notably outperformed the first version of xCell, which ranked tenth with a Spearman rho of 0.724. This improvement highlights the significant advancements made in this updated version. For the fine-grained cell type annotations, xCell 2.0 demonstrated even stronger performance, achieving the highest Spearman correlation (0.666) among all methods. This result surpassed both Aginome-XMU (rho = 0.641) and the first version of xCell (rho = 0.611), underscoring xCell 2.0’s superior ability to capture nuanced cell type differences (Supplementary Figure 3).

A significant challenge in signature-based approaches is the potential for high spillover between closely related cell types. To address this issue, we extended our benchmarking approach to compare spillover levels across the different methods. We quantified spillover by measuring the difference between the correlation of a method’s predictions with the ground truth proportions of the target cell type (direct correlation) and its correlation with the ground truth proportions of the most similar cell type (spill correlation), as described earlier. Using a spillover correction strength (*α*) of 0.5, we found that xCell 2.0 outperformed all other methods in minimizing spillover effects, including compared to the original xCell version (Figure 2D). This superior performance in spillover control, combined with its high accuracy, further emphasizes xCell 2.0’s ability to distinguish between closely related cell types, a crucial factor in accurate cellular deconvolution.

### Prediction of clinical response to immune checkpoint inhibitors

Having established the superior performance of xCell 2.0 in accurately estimating cell type proportions, we sought to apply this method to a clinically relevant challenge in cancer immunotherapy. Immune checkpoint blockade (ICB) therapies have shown remarkable success in treating various cancers, yet patient responses remain heterogeneous. We hypothesized that the composition of the tumor microenvironment (TME) plays a crucial role in determining treatment response. To test this hypothesis, we utilized xCell 2.0 to characterize the TME of 2,007 bulk RNA-seq samples from patients prior to ICB treatment (Figure 3A and Supplementary Table 3). These samples span multiple cancer types and various ICB therapies, including anti-PD-1, anti-CTLA-4, and combination treatments. By applying xCell 2.0 to this diverse dataset, we aimed to identify TME features that could predict patient response to ICB therapy.

**Figure 3.**
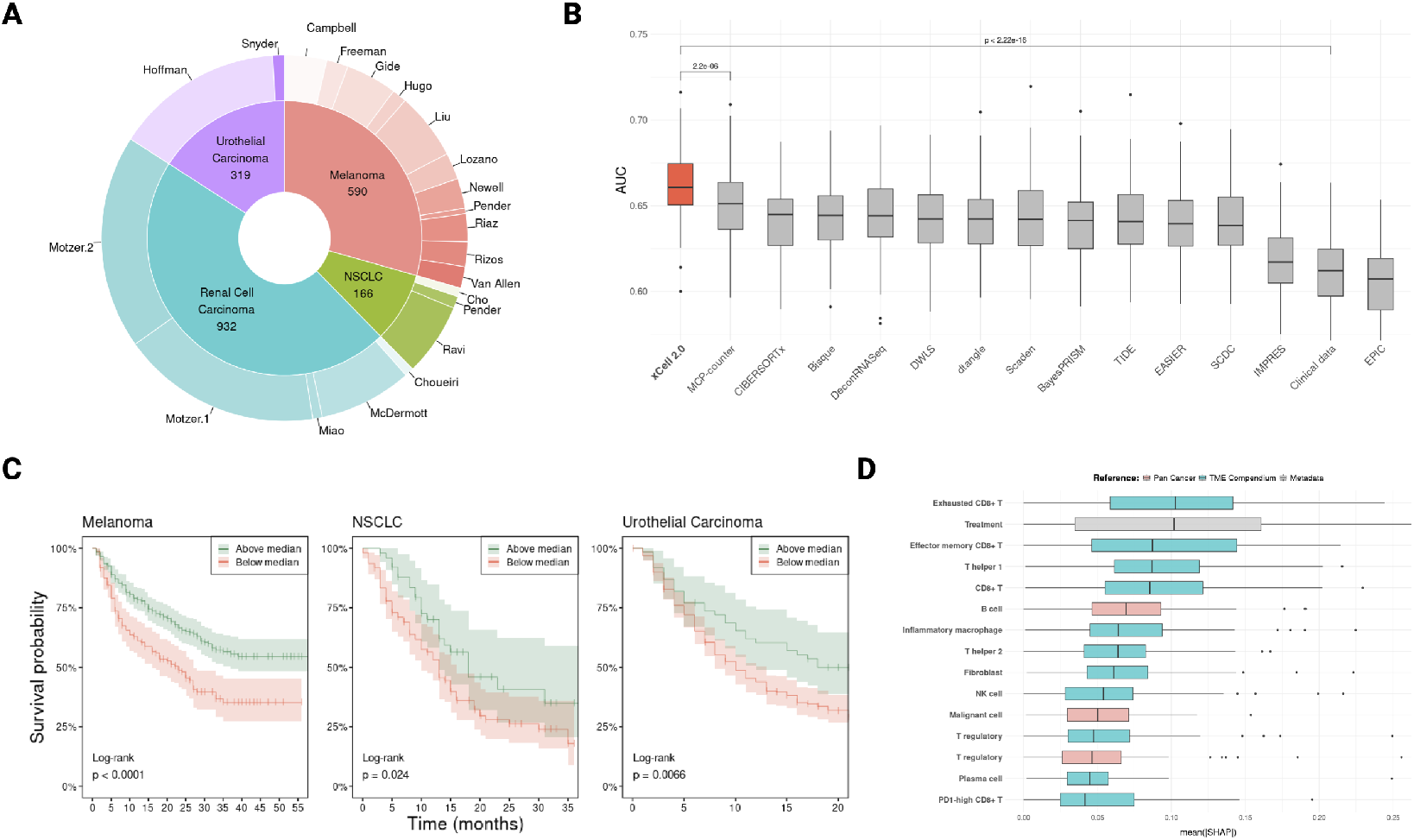
Predicting immune checkpoint blockade response. **A**. Treemap visualization of the datasets used in the ICB response prediction analysis, showing the distribution of samples across different cancer types. **B**. Comparison of prediction performance (AUC) across different deconvolution methods and established ICB response prediction scores. xCell 2.0 demonstrates superior performance (median AUC = 0.661) compared to other methods (p-value *<*2e-16 compared to clinical data alone, p *<*2e-6 compared to the second-best method). **C**. Kaplan-Meier survival curves for patients stratified by predicted ICB response (high 50% vs low 50%) in three cancer types: Non-Small Cell Lung Cancer (NSCLC), Urothelial Carcinoma, and Melanoma. Patients predicted to have high response show significantly better survival outcomes across all three cancer types (NSCLC: p = 0.024, Urothelial Carcinoma: p = 0.0066, Melanoma: p *<*0.0001). **D**. Boxplots show the distribution of absolute mean SHAP values across 100 iterations of the machine-learning model. Each box represents a feature, with higher values indicating greater importance in predicting ICB response. The features are ordered by their median SHAP value, with the most influential predictors at the top.

To analyze this dataset, we applied xCell 2.0 using two pan-cancer references, generating scores for 50 cell types across all 2,007 samples. We classified patients as responders or non-responders based on RECIST criteria (CR/PR vs. SD/PD). To predict treatment response, we implemented a machine learning pipeline using the LightGBM algorithm, employing a nested cross-validation approach with 100 iterations to ensure robust performance estimation.

Our model, based on xCell 2.0’s TME characterization, achieved a median ROC-AUC of 0.661. This performance significantly surpassed a baseline model using only cancer type and treatment information (AUC=0.612; p-value *<*2e-16), underscoring the importance of TME composition in predicting ICB response (Figure 3B). Notably, when compared to other deconvolution methods and established ICB prediction scores (TIDE [33], IMPRES [34]), xCell 2.0-based predictions demonstrated significantly superior performance (compared to second-ranked p-value *<*2e-6). Furthermore, we observed that our pan-cancer response prediction score correlated significantly with survival outcomes across the three cancer types studied with sufficient sample size 3C).

To interpret our model, we utilized SHAP values [35] to identify key predictive features across all iterations (Figure 3D). Notably, exhausted CD8+ T cells emerged as the most important feature in predicting response to ICB therapy. This finding is particularly significant as it aligns with the known biological role of these cells in the TME and their impact on immunotherapy outcomes [36]. We also note that PD-1 high CD8+ T cells appeared among the top predictors, highlighting one of the key advantages of the ability to train on specialized references for specific tasks. This capability allows for the identification of highly specific cell states that are crucial in particular contexts, such as ICB response prediction. This analysis of feature importance provided by xCell 2.0 offers insights into the multifaceted determinants of ICB response, capturing both immune and non-immune factors that influence treatment outcomes.

## DISCUSSION

We present xCell 2.0, an advanced version of the xCell algorithm that introduces a key feature: the ability to train custom reference datasets. To enable this functionality, we incorporated several critical improvements. These include automated handling of cell type dependencies through ontological integration, an improved gene signature selection process, and automatic generation of in-silico simulations for parameter learning and spillover correction. Our comprehensive benchmarking results demonstrate that xCell 2.0 consistently outperforms other popular methods for cellular deconvolution in terms of accuracy and robustness. Notably, xCell 2.0 also shows superior ability to minimize spillover effects between related cell types, a common challenge in deconvolution methods.

The enhancements in xCell 2.0 significantly increase its flexibility and ease of use. The ability to train custom reference datasets allows users to tailor the algorithm to their specific experimental conditions or tissue types. This feature is particularly valuable when analyzing complex tissues like the TME, which contains cell types not typically found in blood. Furthermore, the automated handling of cell type dependencies using ontological integration streamlines the signature generation process, especially for custom references with many cell types. This automation reduces the need for manual intervention and domain expertise, making the tool more accessible to a broader range of researchers.

One of the key advantages of xCell 2.0 that emerged from our comprehensive benchmarking process is its robustness to differences in normalization between the reference dataset and the mixed dataset under analysis. While we invested considerable effort in identifying optimal normalization strategies for each comparative method, ensuring compatibility between reference and validation sets, xCell 2.0 demonstrated a unique flexibility in this regard. This flexibility stems from its approach: it doesn’t require the reference and mixed datasets to be normalized in the same way or even to originate from the same platform. While careful normalization remains crucial for both the reference data (to ensure accurate gene signature learning) and the mixed dataset (for reliable simulations), the decoupling of these normalization requirements represents a substantial practical advantage. This characteristic significantly simplifies the analytical pipeline and broadens the applicability of xCell 2.0 across diverse datasets.

Our application of xCell 2.0 to a large dataset of over 2,007 cancer patients prior to ICB therapy - the largest such dataset analyzed to date - demonstrates its potential for advancing precision medicine. In this analysis, xCell 2.0 outperformed other deconvolution methods in predicting patient response to ICB therapy, showcasing its superior ability to capture relevant cellular information from bulk gene expression data. Importantly, our models based on xCell 2.0 TME characterization significantly outperformed those using only cancer type and treatment information. This finding highlights the critical role of TME composition in determining treatment outcomes, and the importance of accurate cellular deconvolution in cancer research. The superior performance of xCell 2.0 in predicting ICB response compared to established ICB prediction scores, such as TIDE and IMPRES, further highlights its potential as a valuable tool in cancer research and clinical decision-making. By providing a more accurate picture of the cellular composition of tumors and their microenvironment, xCell 2.0 can offer insights into the immune landscape and potential targets for therapy. This could lead to more informed treatment decisions, potentially improving patient outcomes and reducing unnecessary treatments.

One of the key strengths of xCell 2.0 is its consistent performance across different reference types and biological contexts. While other methods often show variable performance depending on the reference used, xCell 2.0 maintains high accuracy across diverse datasets. This robustness is crucial for researchers working with non-standard tissue types or in fields where well-established reference datasets may not be available. The provision of pre-trained reference objects for various human and mouse tissues, including comprehensive immune cell compendiums and pan-cancer datasets, significantly enhances the accessibility and applicability of xCell 2.0. This feature, combined with the flexibility to accommodate custom references, positions xCell 2.0 as a versatile solution for a wide range of research applications, from basic biology to clinical research.

Despite these advancements, it’s important to acknowledge potential limitations. While the reliance on reference datasets is a common limitation for all such methods, xCell 2.0 also faces challenges in distinguishing closely related cell types, particularly in complex tissues. The method’s performance may vary depending on the heterogeneity of the input samples, potentially leading to less accurate results in highly homogeneous datasets. Additionally, the spillover correction, while effective, may not completely eliminate all cross-talk between cell types, especially in cases of extreme imbalance. Furthermore, while xCell 2.0 shows improved performance in fine-grained cell type annotations, its accuracy may decrease for very rare cell populations or subtypes not well-represented in reference datasets. Lastly, the computational requirements for training custom references and analyzing large datasets may be substantial, potentially limiting its use in resource-constrained environments.

In conclusion, xCell 2.0 represents a significant advancement in the field of cellular deconvolution. Its improved performance, flexibility, and robustness make it a powerful tool for understanding the cellular heterogeneity of complex tissues and diseases. The accurate prediction of patient response to ICB therapy demonstrates its potential for advancing precision medicine, not only in cancer but potentially in other diseases as well. As the field of single-cell genomics continues to expand our understanding of cellular diversity, tools like xCell 2.0 will play a crucial role in translating these insights to the analysis of bulk tissue data, bridging the gap between high-resolution single-cell studies and the more widely available bulk transcriptomics data.

## METHODS

### Data Sources

To develop and validate xCell 2.0, we utilized diverse datasets for cell type references and validation. This section describes the sources and processing of these data.

#### Cell type references

We compiled a collection of six human and three mouse gene expression reference datasets of distinct cell types. Those references were categorized into three groups: mixed tissues (general), blood (organ/tissues specific), and disease-associated. The datasets were processed according to the platform from which they were derived: microarray, bulk RNA-Seq and scRNA-Seq. Two microarray-based references: LM22 [18] and ImmGenData [37] underwent Robust Multi-array Average (RMA) normalization. Bulk and scRNA-Seq references were normalized to Transcripts Per Million (TPM) and to Counts Per Million (CPM) accordingly. Additionally, the raw count data for scRNA-seq references were retained where necessary. Annotations for cell types were obtained from the original sources. For bulk gene expression datasets the annotations were based on cell sorting techniques, while annotations for scRNA-seq datasets were based on different computational methods as described in the original study. We identified the corresponding cell type ontology for each of those cell types and used the *xCell2GetLineage* function to generate a cell type dependency file, which was subsequently verified manually. A comprehensive table of all reference datasets, including their sources and key characteristics, is provided in Supplementary Table 1.

#### Validation datasets

To evaluate the performance of xCell 2.0, we created a validation dataset collection compiled from a total of 26 datasets (24 human and 2 mouse). Each validation dataset comprises two matrices: one representing bulk gene expression profiles of cell type mixtures, and the other providing the corresponding ground truth proportions of these cell types. Regarding the ground truth cell type proportions, 22 datasets utilized cytometry techniques (19 flow cytometry and 3 mass cytometry), while the remaining four employed alternative methods such as cell slides, hematology analyzers, and immunofluorescence. The bulk gene expression mixtures were normalized to RMA or TPM in accordance with the specific platform used for gene expression measurement. Detailed information about each validation dataset, including its source, tissue type, and ground-truth methodology, is available in Supplementary Table 2.

### xCell 2.0 Custom Reference Training Pipeline

The xCell 2.0 custom reference training pipeline consists of several key steps designed to generate cell type signatures and learn linear transformation and spillover parameters for accurate estimation of cell type proportions from bulk gene expression data. This section details each step of the pipeline, highlighting improvements over the original xCell method.

#### Pseudo-bulk generation (scRNA-Seq references only)

For scRNA-Seq references, we first generate pseudo-bulk samples by aggregating cell counts. This process is controlled by two parameters: the minimum number of cells per sample (min n cells) and the minimum number of pseudo-bulk samples (min ps samples) per cell type. We calculate the maximum number of possible pseudo-bulk samples given min n cells and the total number of cells available for each cell type in the reference. If the maximum number of samples is less than min ps samples, we adjust min n cells to allow fewer cells per sample. Cells are then randomly selected for each sample per cell type, and their counts are summed for every gene.

#### Preparing input Data

This step involves a few preprocessing procedures for both the input reference and mixture data (if provided). For scRNA-seq references, pseudo-bulk samples are normalized to counts per million (CPM). Following normalization, we select only the most variable genes, ensuring a minimum number of genes for analysis, controlled by the min sc genes parameter. To determine the most variable genes, we calculate the variance of each gene in the reference dataset and iteratively apply the following variance cutoff values: 1.5, 1.0, 0.8, 0.5, 0.3, 0.1, filtering out genes until the remaining gene count exceeds the min sc genes threshold. Next, both bulk and scRNA-seq reference data are converted to log space by adding 3 to restrict inclusion of small changes and followed by log2 conversion. Finally, we identify and retain only the genes shared between the input reference and the mixture data (if applicable). Additionally, an option is provided to convert human gene symbols to their mouse equivalents, enabling cross-species cell type enrichment analysis when required.

#### Identifying cell type hierarchical relationships

A key improvement in xCell 2.0 is the automated handling of cell type dependencies. We use each cell type’s unique ontology ID to consider lineage-dependent cell types in all the following steps. For this purpose, we developed the *xCell2GetLineage* function, which automatically prepares a list for each cell type of its ancestors and descendants present in the reference. This function loads cell type ontology data (updated to 2023) from https://www.ebi.ac.uk/ols4/ontologies/cl using the ontoProc package v1.22.0 [38] and detects lineage relationships using the ontologyIndex package v2.12 [39]. The *xCell2GetLineage* function can be called directly from within the main xCell 2.0 pipeline or used independently to generate a lineage table file that can be manually inspected and provided to xCell 2.0 as an optional argument.

#### Signatures generation

The core methodology and parameters for the signature generation process are consistent with those described in Aran et al. [8], with one key modification. Instead of repeatedly applying the gene inclusion criteria based on the first to third largest quantile expression value of all other cell types, we introduce a parameter called min_frac_ct_passed, set by default to 50%, to adjust the number of cell types considered in the gene inclusion criteria. For instance, if cell type A is compared to ten other cell types, we apply the criteria for the top five cell types quantile expression values.

We also address the following special scenarios where cell types may not generate at least three signatures using the initial parameters:

1. If a cell type fails to produce at least three signatures with min frac ct passed = 0.5, we incrementally relax this parameter by 5% until either at least three signatures are generated or min frac ct passed reaches 5%.
2. If the cell type still does not yield at least three signatures after relaxing min frac ct passed, we further attempt to relax other thresholds of the initial parameters.
3. In cases where certain cell types generate too many differentially expressed genes in each signature, leading to the cancellation of all signatures due to exceeding the maximum allowed genes per signature, we tighten the initial thresholds.

#### In-silico simulations

We generated a set of simulations containing varying fractions of the Cell Type of Interest (CTOI) expression completed with a control cell type expression. Control cell types, chosen for their minimal correlation with the CTOI to mitigate spillover effects, serve to emulate background noise in gene expression simulation. Specifically, for each CTOI, we used the corresponding mean expression vector from the reference matrix and multiplied this vector with fractions spanning from 1% to 25%. A complementary matrix for the control cell type was generated using 1 minus the CTOI’s fraction values and combined with the CTOI matrix to produce the final simulation. We made sure that CTOI and control do not have cell type lineage dependencies. Finally, mean gene-set enrichment scores were computed using singscore v1.20.0 [40] for each cell type simulation using all signatures.

#### Learning linear transformation and spillover parameters

Using the simulations we learn linear transformation parameters for each cell type using the nlsLM function from the minpack.lm package v1.2.4 with the following formula:

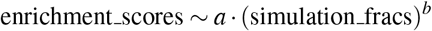

After learning the transformation parameters we fit a linear regression model between the transformed enrichment scores and the simulation fraction to learn the slope and y-intersect for estimating cell type fraction instead of scores. The methodology for learning spillover parameters remains the same as the original xCell method.

#### Reference object construction

The final output of *xCell2Train* is an R S4 object that consists of all cell types’ signatures, a list of cell type dependencies, linear transformation parameters, a spillover matrix, and a vector of the genes used to generate this reference. This object is required as input for cell type enrichment analysis using *xCell2Analysis*.

#### Final cell type proportion estimation

To estimate cell type proportions of the input bulk mixture, we use the *xCell2Analysis* function. In *xCell2Analysis*, we first calculate mean enrichment scores for each cell type by all the corresponding signatures using the singscore v1.20.0 method [40]. These scores are then transformed to fractions using the linear transformation parameters using a regression model and adjusted for spillover using the spillover matrix. The spillover correction strength can be controlled by the user through the parameter *α* (default = 0.5), allowing for fine-tuning of the correction process.

The final output of xCell 2.0 is a matrix of estimated cell type proportions, with rows corresponding to cell types and columns to samples. These estimates represent the relative abundance of each cell type in the input samples, providing a comprehensive view of cellular composition from bulk gene expression data.

### Benchmarking

We conducted a comprehensive benchmarking analysis to evaluate the performance of xCell 2.0 in comparison to other gene-set enrichment and deconvolution methodologies. This comparative assessment was executed across diverse testing scenarios, encompassing various tissue types, data modalities, and validation strategies.

To maximize results accuracy, we implemented a systematic reference-validation matching process where both reference and validation datasets were categorized based on their tissue of origin. Subsequently, we performed dataset matching such that blood/immune and tumor references were exclusively paired with their corresponding validation datasets. General references, which encompass cell types from multiple tissues (e.g., Blueprint-Encode), were matched with all validation datasets. For each matched pair, only cell types common to both the reference and validation datasets were considered, with a minimum threshold of three shared cell types. This yielded a set comprising 85 (79 for human and 6 for mouse) matched pairs between references and validation datasets.

To ensure a fair as possible comparison across all methods, especially those with limited support for custom references and different gene expression measurement platforms, we employed a standardized preprocessing workflow to generate the necessary inputs for each algorithm. This workflow utilized a combination of our own functions and external tools. The specific requirements for each method are detailed below and in Supplementary Table 4. Briefly, we made sure to use input data in the recommended scale (linear or logarithmic) for each method. In addition, if not recommended otherwise, similar normalization was used to all input data if possible. To maximize consistency in normalization across different gene expression platforms, all microarray-based data underwent RMA normalization and scRNA-Seq data normalized to CPM. Bulk RNA-seq references and validations data all normalized in TPM except two validation sets which underwent CPM normalization.

For methods requiring signature and/or gene expression profile matrices and lack built-in functions, we used CIBERSORTx to generate these matrices. We developed a function to preprocess our references and provide all necessary inputs for CIBERSORTx, including the generation of the cell type pheno file, which considers cell type dependencies. The function adheres to recommended settings for different reference types, such as applying quantile normalization for microarray data, using *rmbatchSmode* for scRNA-seq data, or *rmbatchBmode* for bulk reference data. One exception was the LM22 reference which was used in its original form. Additionally, for methods that require cell type marker gene lists, we identified those markers using a function from dtangle which follows the methodology of Abbas et al. [41]. The function is available on dtangle’s GitHub repository.

#### Handling cell type dependencies in deconvolution-based methods

In deconvolution-based methods, the estimated proportions of cell types are interdependent, as the total proportions must sum to one. Thus, estimating the proportions of cell types with lineage dependencies should be avoided. To account for this, we ran multiple iterations of each deconvolution algorithm for the same reference-validation pair. First, we used the *xCell2GetLineage* function to identify lineage relationships among those cell types. These relationships were considered as dependencies, similar to how xCell 2.0 manages such dependencies. Next, we ran each deconvolution algorithm using only the independent cell types from the bottom of the lineage hierarchy. This initial run provided baseline proportions for most of the cell types in the reference. In subsequent runs, we systematically excluded descendant cell types from each lineage hierarchy and replaced them with the corresponding parent cell type. For example, if the reference-validation pair included T cells, CD4+ T cells, CD4+ naïve T cells and CD4+ memory T cells, the initial run would include CD4+ naïve T cells and CD4+ memory T cells. The second run would exclude those cell types and include only CD4+ T cells. Finally, the third run would include only general T cells. After completing all iterations, we combined the proportions from the initial run with those of the dependent cell types to create a final proportions matrix. This procedure extended the run time of the deconvolution-based methods significantly.

#### Performance evaluation

To ensure a fair and robust comparison of method performance across datasets of varying sizes, we implemented a weighted correlation analysis approach. For each method and dataset combination, we calculated both Spearman and Pearson correlation coefficients between the predicted and ground-truth cell type proportions. To account for the disparity in sample sizes across validation datasets, we applied the following weighting procedure:

1. Fisher’s z-transformation was applied to the correlation coefficients:

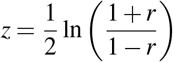

where *r* is the correlation coefficient.
2. The z-scores were then weighted by the natural logarithm of the number of samples in each dataset. To prevent over-representation of large datasets, we capped the effective sample size at 30. For datasets with more than 30 samples, we used 30 as the weighting factor. This can be expressed as:

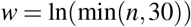

where *n* is the number of samples in the dataset.
3. The weighted z-scores were calculated as:

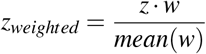
4. Finally, we applied the inverse Fisher transformation to convert the weighted z-scores back to correlation coefficients:

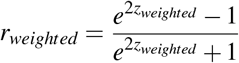

This approach ensures that larger datasets contribute more to the overall performance assessment while preventing them from dominating the results. It also maintains the interpretability of the final metrics as correlation coefficients, allowing for intuitive comparison across methods and datasets.

#### Cellular composition analysis methods

##### dtangle

We transformed both bulk mixtures and references into log-space as recommended by Hunt et al. [26]. Following dtangle’s vignette, we first normalized scRNA-seq based references to CPM, then transformed them to log-space and applied quantile normalization to both reference and mixture data using the *normalizeBetweenArrays* function from limma [42]. For bulk references, we set marker method to “p.value”, while for scRNA-seq references, we set it to “ratio”. The data type parameter was set to the appropriate reference data type (“rna-seq” or “microarray-gene”). Version 2.0.9 was used.

##### EPIC

Linear scale expression levels were used, as recommended by the developers. If possible, all input data were normalized similarly, ideally to TPM. For custom reference preparation, EPIC requires signature genes, cell type Gene Expression Profiles (GEPs), and optionally, cell type gene variability. Since EPIC lacks built-in functions for identifying signature genes and generating GEP matrices, we used the signature matrix genes and GEP matrix from CIBERSORTx. Gene variability was calculated as the standard deviation of each gene in the original reference. When running EPIC, the withOtherCells parameter was set to FALSE. Although EPIC was not specifically designed for scRNA-seq references, we followed the authors’ suggestions on GitHub for using such references. Version 1.1.6 was used.

##### BayesPrism

Linear scale expression levels were used. The developers recommend using non-normalized count data, though linear normalization (e.g., TPM, RPM, RPKM, FPKM) is acceptable. Following BayesPrism’s vignette, we used the *cleanup.genes* function with parameters: species=“hs” or “mm” (depending on the validation data organism) and gene.group=c(“Rb”,”Mrp”, “other Rb”, “chrM”, “MALAT1”, “chrX”, “chrY”). We selected protein-coding genes using the select.gene.type function. As BayesPrism only supports marker gene identification for scRNA-Seq references, we identified marker genes for bulk references using dtangle. We created a prism object using *new.prism*, setting the input.type parameter to “count.matrix” for scRNA-seq references and “GEP” for bulk references, specifying the key for malignant cell labels, and running BayesPrism with 45 CPUs using *run.prism*. Version 2.1.1 was used.

##### MCP-counter

This is an enrichment scores-based method, requiring only a set of gene signatures per cell type. We used the *MCPcounter.estimate* function with the genes parameter set to a data frame containing all marker (signature) genes for each cell type, identified using dtangle. Version v1.2.0 was used.

##### DeconRNASeq

Linear scale expression levels were used. The developers recommend using the same normalization for all input data. DeconRNASeq requires a signatures matrix but does not provide a built-in function to generate it. We therefore used the signature matrix generated by CIBERSORTx. Version 1.42.0 was used.

##### CIBERSORTx

This tool provides methods for handling different gene expression platforms and all necessary functions to generate required inputs. Details about generating signatures and GEP matrices using CIBERSORTx are mentioned above.

##### Bisque

We used the implementation from the omnideconv package v0.1.0 [43] using the *deconvolute_bisque* function. Reference and bulk mixture data were provided in linear space. For bulk references comprising multiple datasets or scRNA-seq references from different individuals, we provided this information to the batch_ids argument. We also used the “markers” argument of *deconvolute_bisque* with cell type markers identified by dtangle.

##### DWLS

Linear scale expression levels were used. Due to inconsistencies in the built-in signature generation function across all references, we used the signature matrix generated by CIBERSORTx. Deconvolution was performed using the *solveSVR* function with default parameters. Version 0.1.0 was used.

##### SCDC

The method was implemented using the omnideconv package [43], specifically the *deconvolute_scdc* function. Information regarding different datasets or individuals in the reference was provided through the *batch_ids* parameter. Input data was used in linear scale.

##### Scaden

The method was implemented using the omnideconv package [43] via the *deconvolute scaden* function, which runs Scaden in Python through R. The model was generated using the *build_model_scaden* function, and cell type proportions were obtained using the *deconvolute_scaden* function. We followed the default parameters as recommended in the omnideconv package documentation.

##### DREAM challenge assessment

We evaluated xCell 2.0 using the data and instructions provided in the run-deconvolution-method-on-challenge-data. R script available on the GitHub repository associated with the study by White et al. [32]. To comprehensively assess all cell type annotations included in the challenge data, both at the coarse and fine-grained levels, we combined three bulk RNA-Seq reference datasets — BluePrint-Encode [14, 15, 44], Blood Immune Compendium [17], and Database of Immune Cell Expression (DICE) [45] — to create an xCell 2.0 object specifically tailored for this evaluation.

### Machine Learning Analysis for ICB Response Prediction

To demonstrate the practical application of xCell 2.0, we trained a machine learning model to predict patient response to immune checkpoint blockade (ICB) therapy based on tumor microenvironment (TME) composition.

#### Dataset Preparation

We curated a comprehensive pan-cancer dataset comprising bulk RNA-seq data from 2,007 patients across 20 datasets, representing the largest collection of its kind to date for ICB treatment analysis. This dataset includes samples from multiple solid tumor types, including melanoma, renal cell carcinoma (RCC), non-small cell lung cancer (NSCLC), and urothelial cancer (UC), treated with various ICB therapies such as anti-PD-1, anti-CTLA-4, and combination treatments.

Data were collected from publicly accessible resources, including supplementary tables from research papers and the Gene Expression Omnibus (GEO). Patient response was classified based on RECIST criteria, with complete response (CR) and partial response (PR) grouped as responders, and stable disease (SD) and progressive disease (PD) as non-responders. For one large dataset (354 samples) lacking RECIST values, overall response data was used.

The datasets were merged based on shared genes, resulting in approximately 15,000 common genes across all samples. Data normalization (TPM) was performed to ensure comparability across datasets. Transcript lengths were obtained using the biomaRt package v2.58.0 in R, focusing on canonical transcripts. All studies used to compose this dataset are detailed in Supplementary Table 3. Importantly, since xCell 2.0 primarily utilizes the ranking order of gene expression, our analysis did not require special normalization of the integrated dataset beyond the procedures described above.

#### Feature Generation

We applied xCell 2.0, CIBERSORTx, dtangle, EPIC, MCP-counter, BayesPrism, DeconRNASeq, Bisque, DWLS, SCDC and Scaden to the curated dataset using two pan-cancer reference objects. Using the same methodology described in the benchmarking section, each method results in the cell type proportion estimates for 40 cell types (10 were shared between the references). These cell type proportions served as features for our prediction model. Additionally, we included cancer type and treatment information as baseline features.

#### Model pipeline

We employed the LightGBM algorithm [46] using the R package lightgbm v4.4.0, a gradient boosting framework that uses tree-based learning algorithms, for our prediction model. To ensure robust performance estimation and mitigate overfitting, we implemented a nested cross-validation approach, with extensive hyperparameter tuning. For each iteration, the dataset was split into training (75%) and testing (25%) sets using stratified sampling to maintain the distribution of responders and non-responders. The RECIST response variable was binarized, with complete response (CR) and partial response (PR) coded as 1 (responders), and stable disease (SD) and progressive disease (PD) coded as 0 (non-responders). The entire process was repeated 100 times with different random seeds to account for variability in data splitting.

#### Hyperparameter tuning

We used a multi-staged hyperparamter tuning approach. First, we optimized the learning rate through an initial search over a predefined range, followed by a refinement step around the best initial rate. This refinement explored a range of ±80% around the initial best value, using 10-fold cross-validation for each candidate rate. The learning rate yielding the highest Area Under the ROC Curve (AUC) was selected for subsequent stages. Next, we conducted two stages of grid search to fine-tune other crucial hyperparameters. The first stage focused on the model structure, exploring combinations of the number of leaves (as a factor of maximum tree depth), minimum data in leaf, minimum gain to split, and maximum tree depth. The second stage delved into regularization and sampling parameters, tuning the feature fraction, bagging fraction, and L1 and L2 regularization terms.

The final LightGBM model incorporated the best hyperparameters found during this extensive tuning process. We employed binary classification as the objective, with AUC as the primary evaluation metric. The model utilized the GBDT (Gradient Boosting Decision Tree). We enabled early stopping with a patience of 10 rounds and allowed up to 1000 iterations, subject to the early stopping criterion.

#### Model evaluation

We evaluated our model’s performance using the AUC-ROC metric on the held-out test sets from the outer cross-validation loop. We compared the performance of three types of models: (1) Baseline model, using only cancer type and treatment information; (2) Models for xCell 2.0 and other deconvolution methods using TME features; (3) Established ICB prediction scores (TIDE [33], IMPRES [34]). Statistical significance of performance differences was assessed using paired t-tests on the AUC-ROC values across the 100 iterations.

#### Feature importance analysis

To interpret the contributions of different cell types to the prediction, we employed SHAP (SHapley Additive exPlanations) values [35]. SHAP values were calculated for each feature across all iterations of the cross-validation process. We then aggregated these values to identify the most consistently important features for predicting ICB response.

#### Survival analysis

To further validate the clinical relevance of our predictions, we performed a survival analysis. We stratified patients into high- and low-response probability groups based on our model’s predictions and compared their survival outcomes using Kaplan-Meier curves and log-rank tests. This analysis was conducted separately for each cancer type to account for differences in baseline survival rates.

### Code Availability

To ensure reproducibility and facilitate widespread use of xCell 2.0, we have made the codebase and associated resources publicly available in link - …

#### xCell 2.0 package

The xCell 2.0 algorithm has been implemented as an R package, which will be available through the Bioconductor repository (currently in https://github.com/AlmogAngel/xCell2). The package includes the core functions for running xCell 2.0, as well as utility functions for data preprocessing, visualization, and result interpretation. Detailed documentation and vignettes are provided to guide users through the installation process and typical workflows.

#### Pre-trained reference objects

We have made available a comprehensive collection of pre-trained reference objects for both human and mouse tissues. These objects, which include the signature matrices, transformation models, and spillover parameters, can be downloaded from our dedicated repository (https://dviraran.github.io/xCell2refs). This resource allows users to immediately apply xCell 2.0 to their datasets without the need for custom reference training.

#### Web application

To increase accessibility for researchers without programming experience, we have developed a web-based interface for xCell 2.0 (https://aranlab.shinyapps.io/xCell2/). This application allows users to upload their gene expression data and obtain cell type proportion estimates using our pre-trained references. The web tool also provides basic visualization and data export capabilities.

#### Benchmark and analysis scripts

The scripts used for benchmarking xCell 2.0 against other methods and for the ICB response prediction analysis are available on GitHub (https://github.com/AlmogAngel/xCell2). This repository includes code for data preprocessing, running comparisons with other deconvolution methods, and implementing the machine learning pipeline for ICB response prediction.

#### License

xCell 2.0 is released under the GNU General Public License (GPL) v3.0.

## Acknowledgments

Research in the Aran lab is supported by grant from the Israel Science Foundation (1543/21). DA is also supported by the Azrieli Faculty Fellowship. We thank the Aran Lab members for helpful comments.

## Author Contributions

AA developed the method and performed all benchmarking analyses. LN collected the immunotherapy data and performed the related analyses. SNL and AA supported the immunotherapy analysis. DA supervised all aspects of the study and wrote the manuscript with support from AA. All authors reviewed and approved the final manuscript.

