## Supplementary Figures for "xCell 2.0: Robust Algorithm for cell type Proportion Estimation Predicts Response to Immune Checkpoint Blockade"

### Supplementary Figure 1

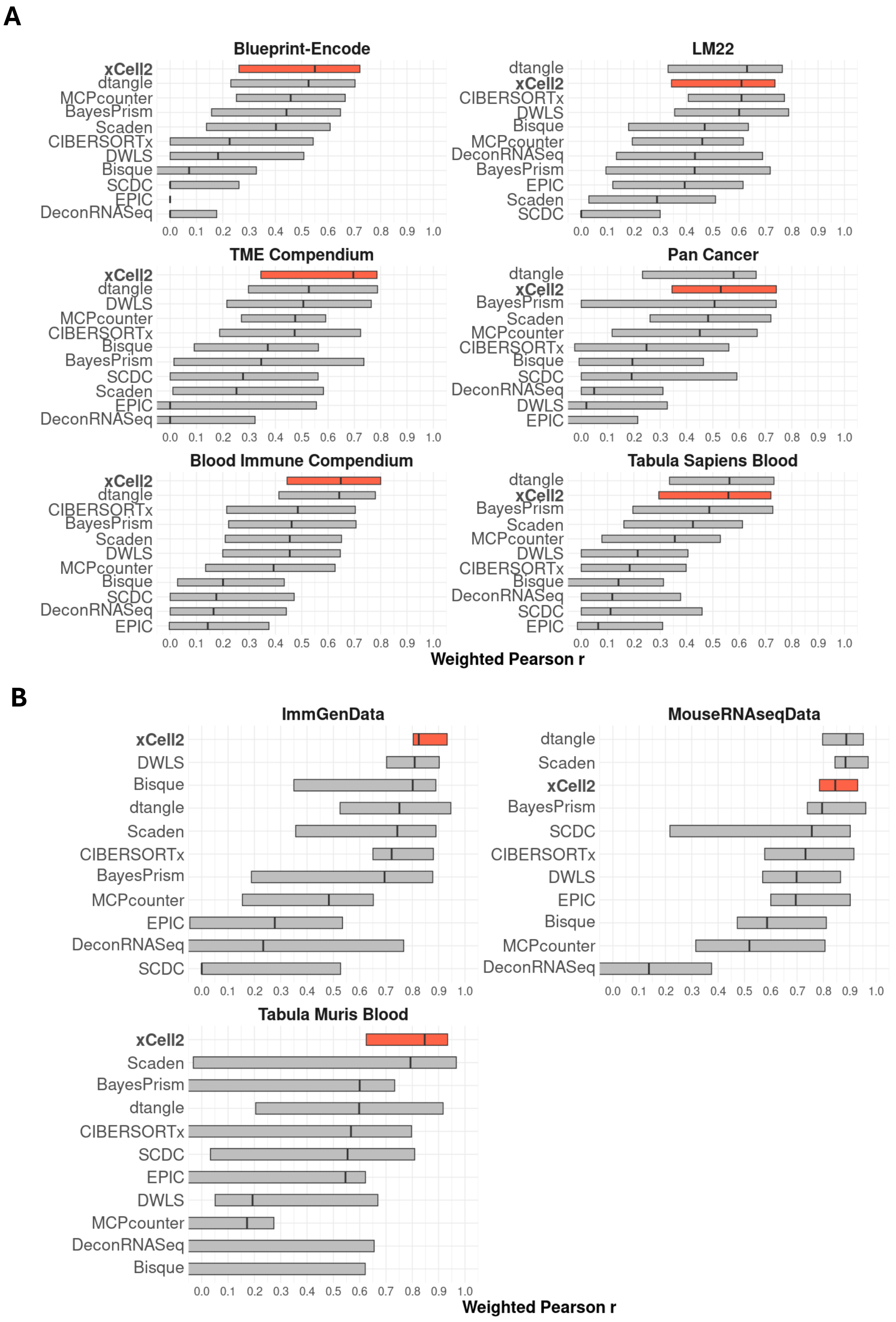


**Supplementary Figure 1.** Performance benchmarking of xCell 2.0 against other deconvolution methods displayed by method and reference in weighted Pearson correlation (r) for multiple human (A) and mouse (B) reference datasets.

### Supplementary Figure 2

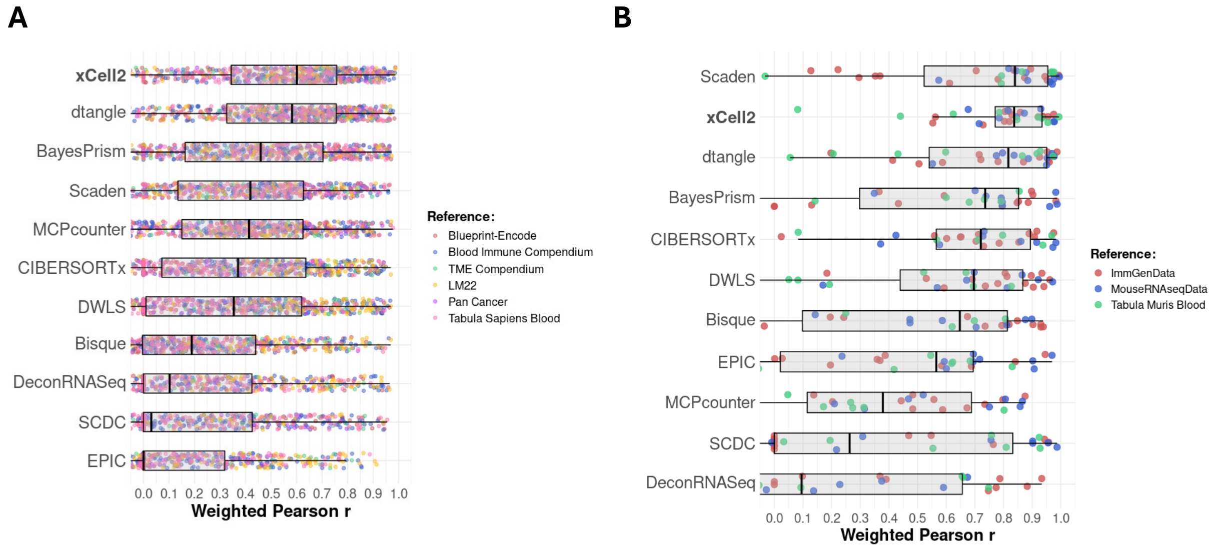


**Supplementary Figure 2.** Performance benchmarking of xCell 2.0 against other deconvolution methods displayed by method in weighted Pearson correlation (r) for multiple human (A) and mouse (B) reference datasets.

### Supplementary Figure 3

**A**

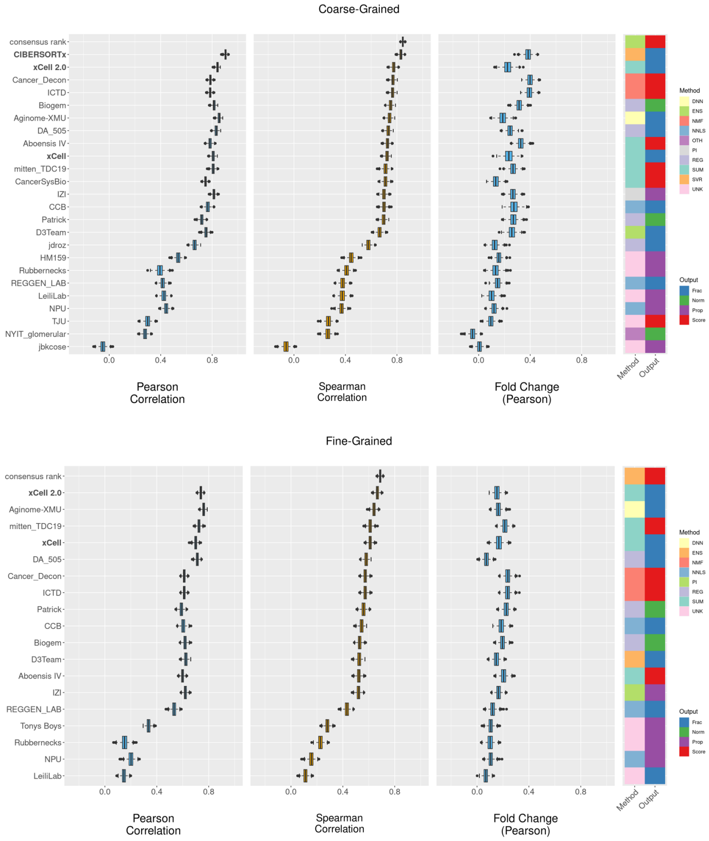


**B**

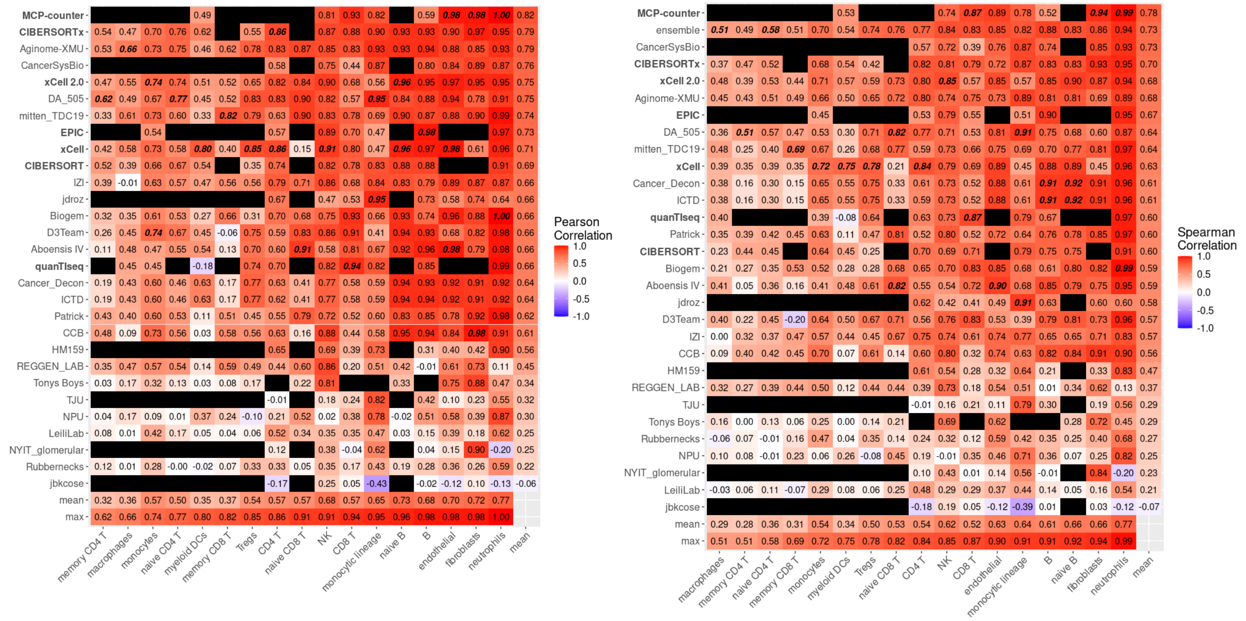


**Supplementary Figure 3.** Performance comparison of xCell 2.0 with other deconvolution methods in the DREAM Challenge datasets. Boxplots (A-B) and heatmaps (B, C) replicate the figures from White et al. (2024) with the addition of xCell 2.0.
